# The human immune checkpoint molecule, HLA-G2, induces tolerance in monocytes and dendritic cells via upregulation of PD-L1

**DOI:** 10.1101/2023.10.14.562373

**Authors:** Ami Takahashi, Kimiko Kuroki, Naoyoshi Maeda, Mie Nieda, Katsumi Maenaka

## Abstract

Human leukocyte antigen (HLA)-G is a non-classical HLA class I immunomodulatory molecule with restricted expression in the placenta, thymus and regulatory T cells. The spliced isoforms of HLA-G include an α2 domain-deleted isoform, HLA-G2, which specifically binds to the immune checkpoint leukocyte immunoglobulin-like receptor B2 (LILRB2), to suppress immune responses in myelomonocytic cells. We previously reported the structural and receptor binding characteristics of recombinant HLA-G2 protein and its immunosuppressive effects on inflammation in mouse models. However, the function and the mechanism of action of HLA-G2 on human immune cells have not been elucidated.

In the present study, we demonstrate the immunosuppressive effect of HLA-G2 on human CD14-positive monocytic cells. HLA-G2 induced the production of the immunosuppressive cytokine, IL-10, and stimulated IL-6/STAT3/indoleamine-2,3-dioxygenase signaling by binding to LILRB2. HLA-G2 binding to LILRB2 also down-regulated cell surface expression of HLA-DR and CD86. Unexpectedly, HLA-G2 up-regulated cell surface expression of PD-L1 in both CD14-positive monocytic cells and interferon-induced dendritic cells (IFN-DCs). This observation suggests HLA-G2/LILRB2 signaling promotes PD-L1 expression. Furthermore, HLA-G2 treatment of IFN-DCs suppressed T cell proliferation in mixed lymphocyte reactions. These findings provide novel insights into the modulation of human immune responses of tolerogenic myelomonocytic cells induced by HLA-G2 binding to LILRB2, and suggest that targeting the HLA-G2-LILRB2 interaction could be a novel approach for immune checkpoint therapy.

**Significance statement:** During pregnancy, HLA-G isoforms are expressed by fetal trophoblasts to suppress maternal immune responses. Among various HLA-G isoforms, the HLA-G2 homodimer has been expected as an immunosuppressive biologic targeting myelomonocytic antigen-presenting cells via leukocyte immunoglobulin-like receptor B2. We previously reported significant immunosuppressive effects of HLA-G2 in autoimmune mouse models. Here, we first demonstrate that HLA-G2 isoform induces tolerogenic phenotypes of human peripheral immune cells by significantly upregulating an immune checkpoint molecule, PD-L1. Monocyte-derived dendritic cells stimulated by HLA-G2 suppressed T cell proliferation in mixed lymphocyte reactions. These results suggest that HLA-G2 can be a novel candidate for immune checkpoint therapy.

## Introduction

Human leukocyte antigen-G (HLA-G) plays an important role in immune tolerance. The characteristics of HLA-G are distinct from classical HLA class I molecules: low polymorphism, restricted tissue distribution, and expression as several spliced isoforms (1–3). The restricted expression of HLA-G on placental trophoblasts contributes to maternal-fetal immune tolerance (4). HLA-G is also expressed in thymic tissue (5) and regulatory T cells (6). HLA-G expression is also observed in pathological conditions such as tumorigenesis, viral infections, and transplantation (7). To date, seven spliced isoforms of HLA-G have been reported. HLA-G1 to HLA-G4 are membrane-bound isoforms and HLA-G5 to HLA-G7 are soluble isoforms (1–3) (Supplementary Figure 1). HLA-G1 and HLA-G5, which have identical extracellular domains, share a similar structural organization to classical HLA class I molecules composed of a heavy chain (α1, α2 and α3 domains), a peptide, and β2-microglobulin (β2m). By contrast, HLA-G2 and HLA-G6, which have identical extracellular domains, form a non-covalently linked homodimer of two α2 domain-deleted heavy chains (α1 and α3 domains), which structurally resembles the heterodimer of HLA class II molecules (8). Hereafter, HLA-G2 refers to the ectodomain of HLA-G2 and HLA-G6 (Supplementary Figure 1).

HLA-G isoforms inhibit immune responses by binding the inhibitory leukocyte immunoglobulin (Ig)-like receptors (LILR) B1 (Ig-like transcript (ILT2)) and LILRB2 (ILT4). LILRB1 and LILRB2 transmit inhibitory signals to immune cells by binding broadly to classical and non-classical HLA class I molecule under physiological conditions. The binding affinity of HLA-G1 to LILRB receptors is higher than other HLA class I molecules (9). LILRB1 is expressed on a broad range of immune cells such as antigen presenting cells (APCs), T cells and natural killer (NK) cells. By contrast, LILRB2 is mainly expressed on myelomonocytic cells, including monocytes, macrophages, and dendritic cells (DCs) (10). A mouse orthologue of human LILRB2, the paired Ig-like receptor (PIR)-B, is expressed on APCs and binds to both HLA-G1 and HLA-G2 (11, 12). Both HLA-G1 and HLA-G2 promote the immunosuppression in inflammatory disease mouse models (11–16).

The immunosuppressive mechanism of HLA-G1 has been widely studied. HLA-G1 binding to LILRB1 broadly inhibits immune cell function, such as proliferation and differentiation of T and B cells, and Ig secretion (17, 18). In addition, HLA-G1 inhibits DC maturation in part by downregulating HLA class II, CD80, and CD86, and prolongs the allograft survival (19, 20). HLA-G1 binding to LILRB2 modulates the IL-6 signaling pathway and STAT3 activation in LILRB2 transgenic mice and human monocyte-derived DCs (19, 21). In human macrophages, soluble HLA-G1 increases IL-6 and indoleamine-2,3-dioxygenase 1 (IDO-1) production, which inhibits T cell activation (22). By contrast to HLA-G1, the function and mechanism of action of the HLA-G2 isoform remains unclear.

Individuals homozygous for the *HLA-G*0105N* allele do not express the functional HLA-G1 isoform but can still produce α2 domain-deleted HLA-G2 and HLA-G6 isoforms. Interestingly, HLA-G2 bound to LILRB2 with high affinity by avidity effect (8). Furthermore, the soluble HLA-G2 protein showed anti-inflammatory effects in a rheumatoid arthritis (RA) mouse model at lower doses compared to HLA-G1 monomers and homodimers (11, 12). These observations clearly support the functional significance of HLA-G2 in immune suppression *in vivo*. However, the effects of HLA-G2 on signaling in human cells have not been defined.

In this study, we investigated the functional effect of HLA-G2 on human LILRB2-expressing CD14-positive monocytes and monocyte-derived dendritic cells. HLA-G2 binding induced an immunosuppressive phenotype in monocytes, eliciting upregulation of two key immunosuppressive molecules programmed death-ligand 1 (PD-L1) and IDO. HLA-G2-treated DCs differentiated from peripheral blood monocytes with GM-CSF and IFN-γ, designated as IFN-DCs (23), also preliminary showed an immunosuppressive phenotype and inhibited proliferation of both CD4-positive and CD8-positive T cells in allogeneic and autologous mixed lymphocyte reactions (MLR), respectively. This study provides a novel insight into the mechanism of the immunosuppressive effects of the unique HLA-G isoform, HLA-G2. Furthermore, it also suggests that targeting HLA-G2−LILRB2 signaling could provide a novel therapeutic intervention in diverse immune disorders.

## Materials and methods

### Preparation of the recombinant HLA-G2 protein

The extracellular domains of HLA-G2 (HLA-G2) were expressed in *Escherichia coli* ClearColi BL21(DE3) competent cells (Lucigen) to avoid triggering endotoxic responses in human cells. The inclusion bodies of HLA-G2 were refolded as previously described (12). Refolded HLA-G2 protein was purified by size exclusion chromatography using HiLoad 26/60 Superdex75 pg column (GE Healthcare) equilibrated with 20 mM Tris-HCl pH 8.0, 100 mM NaCl. The peak of properly refolded HLA-G2 was collected and buffer exchanged into HBS-EP (0.01 M HEPES pH 7.4, 0.15 M NaCl, 3 mM EDTA and 0.005% [v/v] Surfactant P20, GE Healthcare) for surface plasmon resonance (SPR) analysis or PBS for cell-based assays by dialysis. HLA-G2 proteins in PBS were concentrated by Amicon Ultra-15 (NMWL 10,000 Da) (Merck Millipore), and the flowthrough solution from protein concentration was used as a negative control (PBS) in cell-based assays. HLA-G2 and PBS solutions were filter sterilized through a 0.22 μm filter.

### Preparation of human monocytes and DCs

CD14-positive monocytes were isolated from peripheral blood mononuclear cells (PBMCs) with CD14 microbeads and MACS LS column system (Miltenyi Biotec). PBMCs were obtained from the buffy coats of twelve healthy blood donors (eight males and four females, over twenty years old, Hokkaido University) by density gradient centrifugation (Lymphoprep, Veritas) following the manufacturer’s instructions. The mean number of PBMCs was 7.3 ± 1.5 x 10^7^ cells/mL, and CD14-positive monocytes isolated from the PBMCs were 8.2 ± 2.7 x 10^6^ cells/mL. The average purity of CD14-positive monocytes was calculated as 86 ± 7.0% by flow cytometry. CD14-positive cells (1.0 ∼ 2.0 x 10^6^ cells) were incubated with 10% (v/v) HLA-G2 (final concentration of 0.58, 1.2 or 2.3 μM) or 10% (v/v) PBS in the culture medium (RPMI-1640 medium (Wako) containing 10%(v/v) heat-inactivated FBS (Lot no. 10777, Biosera) and Penicillin-Streptomycin-Amphotericin B Suspension (Wako)) at 37°C under 5% CO_2_ condition either overnight or for two days (18 or 48 hours, respectively). Cells cultured in medium alone were used as a non-treated control. CD14-negative lymphocytes were collected for mixed lymphocyte reaction (MLR) assays.

IFN-DCs were differentiated from adherent PBMC cells in 10% FBS RPMI-1640 (Wako) or Gibco AIM V medium (Thermo Fisher Scientific) containing 1000 U/mL GM-CSF (Primmune Inc.) and IFN-α (IFN-α 2b, OriGene Technologies or MSD) as previously described (23). After three days, IFN-DCs were harvested, and cultured in 10% FBS RPMI-1640 or Gibco AIM V medium with 10% (v/v) HLA-G2 (final concentration of 2.3 μM) or 10% (v/v) PBS for two days. Penicillin-Streptomycin-Amphotericin B Suspension (Wako) were added in RPMI-1640 medium.

To generate immature IL-4-DCs, purified CD14-positive cells from PBMCs were cultured in differentiation medium (10% FBS RPMI-1640 (Wako) with 1000 U/mL GM-CSF (Primmune Inc.), 500 U/mL IL-4 (Primmune Inc.) and Penicillin-Streptomycin-Amphotericin B Suspension (Wako)) for seven days.

The clinical study protocol was approved by the Bioethics Committee of Faculty of Pharmaceutical Sciences, Hokkaido University (clinical research no. 2016-006). Informed consents were obtained from all participating healthy donors prior to their inclusion in this study.

### Flow cytometry (FCM) analysis

Cells were harvested and suspended in 50 μL of FCM buffer (0.5% BSA and 0.05% sodium azide in PBS) with 2.5-10 μg of Human BD Fc Block (BD). Cells were stained using the antibodies listed in **Supplementary Table 1** for 15 minutes in the dark at room temperature and washed three times with FCM buffer. Finally, cells were resuspended in 200 μL of FCM buffer including 20 μL 7-amino-actinomycin D (7-AAD) viability staining solution (BD) and incubated for 10-30 minutes before acquisition. A BD FACSCalibur cell analyzer and BD CellQuest version 6.0 software (BD) were used for FCM analysis. Gates for cells staining with antibody were set so that isotype control staining was less than 5% of events. The differences were statistically analyzed using Student’s t-test, and *P* < 0.016 was considered statistically significant by Bonferroni correction,

### Enzyme-Linked Immunosorbent Assay (ELISA)

To evaluate the cytokine levels in cell culture supernatants, ELISA for human IL-6 and IL-10 were performed using ELISA kits (DuoSet [R&D Systems] or BD OptEIA [BD]) following the manufacturer’s instructions. Absorbance at 450 nm (subtracted absorbance at 540 nm) was detected using a Spark 10M (TECAN). The assay ranges for the kits are: IL-6 (9.38-600 pg/mL [DuoSet], 4.7-300 pg/mL [OptEIA]) and IL-10 (31.3-2000 pg/mL [DuoSet], 7.8-500 pg/mL [OptEIA]).

### Western blotting

Cells (2 x 10^6^ cells) were harvested by centrifugation, washed with PBS, and lysed in RIPA buffer (50 mM Tris-HCl pH 8.0, 150 mM NaCl, 2 mM EDTA, 1% Nonidet P-40 [NP-40], 0.5% sodium deoxycholate, 0.1% SDS, cOmplete, EDTA-free protease inhibitor cocktail (Roche), and phosphatase inhibitor cocktail solution I (Wako)). After removal of nuclear pellets by centrifugation, cell lysate supernatants were mixed with sample buffer (final 62.5 mM Tris-HCl pH 6.8, 25% (v/v) glycerol, 2.5% (w/v) SDS, 0.0125% (w/v) bromophenol blue, and 5% 2-mercaptoethanol (2-ME)), separated on the 12.5% or 10% acrylamide gel, and transferred to PVDF membrane (Bio-Rad). Antibodies used in this study are shown in **Supplementary Table 1**. ECL prime western blotting detection reagent (GE Healthcare) and ImageQuant LAS4000mini (GE Healthcare) were used for detection. IDO-1, STAT3, phosphorylated STAT3 and β-actin (as a loading control) were detected on the same membrane by reprobing with antibodies after stripping with stripping buffer (62.5 mM Tris-HCl pH 6.8, 2% SDS, and 0.7% 2-ME).

### Surface plasmon resonance (SPR)

SPR analysis was performed using a BIAcore2000 (GE Healthcare) at 25°C. C-terminally biotinylated LILRB2 and BSA (negative control, Wako) were prepared as previously described (24). Biotinylated proteins were immobilized on a CAP chip (Biotin CAPture kit, GE Healthcare). As a blocking control anti-LILRB2 antibody, clone 27D6 (IgM, Thermo Fisher Scientific) was repeatedly injected until binding to immobilized LILRB2 reached saturation. Then, purified HLA-G2 (3.6 μM) in HBS-EP was injected over either LILRB2, LILRB2 blocked with 27D6 antibody, or BSA (negative control) for 3 min. The flow rate was 10 μL/min.

### Blocking assay

CD14-positive monocytes isolated from human PBMCs (1 x 10^6^ cells/well) were incubated with 27D6 antibody or isotype-matched control antibody (4 μg/mL, IgM, Thermo Fisher Scientific) for 30 min at room temperature, then cultured in 10% FBS RPMI-1640 with 2.3 μM HLA-G2 or PBS for two days. IDO-1 and β-actin (loading control) in the cell lysates were detected by western blotting, and IL-6 and IL-10 in the cell culture supernatants were evaluated by ELISA as described above.

### Allogeneic Mixed Lymphocyte Reaction (MLR)

IFN-DCs incubated with 2.3 μM HLA-G2 or PBS for 48 hours were treated with 10 μg/mL of mitomycin C (Nacalai tesque). CD14-negative PBMC lymphocytes isolated from a different healthy donor were labelled with CFSE to monitor proliferation. CFSE-labelled lymphocytes were co-cultured with IFN-DCs for two days with 100 U/mL IL-2 at a ratio of 10:1 (2×10^6^ cells lymphocytes: 2×10^5^ cells IFN-DC). After 48 hours, proliferating CD3-positive cells were evaluated by FCM using carboxyfluorescein diacetate succinimidyl ester (CFSE) as a proliferation marker.

### Autologous MLR

IFN-DCs generated from a HLA-A*0201-positive donor were stimulated with 2.3 μM HLA-G2 or PBS for two days, and cultured overnight with a Melan-A/MART-1_26-35_ antigenic peptide analogue incorporating an Ala27 to Leu mutation (MART-1(A27L), ELAGIGILTV) (25–27). Subsequently, IFN-DCs were harvested and incubated with lymphocytes from the same donor in RPMI-1640 medium with 10% FBS and IL-2 (100 U/mL) at a 10:1 ratio (2 x 10^6^ cells lymphocytes: 2 x 10^5^ IFN-DCs). After 14 days, CD8-positive MART-1(A27L) peptide-specific T cells were evaluated by FCM by staining with PE-conjugated MART-1(A27L) tetramer and anti-CD8 FITC (BioLegend) (**Supplementary Table 1**).

## Results

### HLA-G2 induced an immunosuppressive phenotype in human monocytes

Recombinant HLA-G2 homodimer protein was prepared as previously described (**Supplementary Figure 2**) (8, 12). LILRB2, a functional receptor for HLA-G2, was constitutively expressed on CD14-positive monocytes isolated from PBMCs (82 ± 16%, *n* = 8) (**Figure 1A**). First, CD14-positive monocytes were treated with different concentrations of HLA-G2 recombinant protein to determine the effective dose of HLA-G2. Production of IL-6, a cytokine induced by HLA-G1 binding to LILRB2 (21), was dose-dependently upregulated by HLA-G2 stimulation (**Supplementary Figure 3B**). Therefore, we evaluated the effects of HLA-G2 at the concentration of 2.3 μM in the present study.

**Figure 1.**
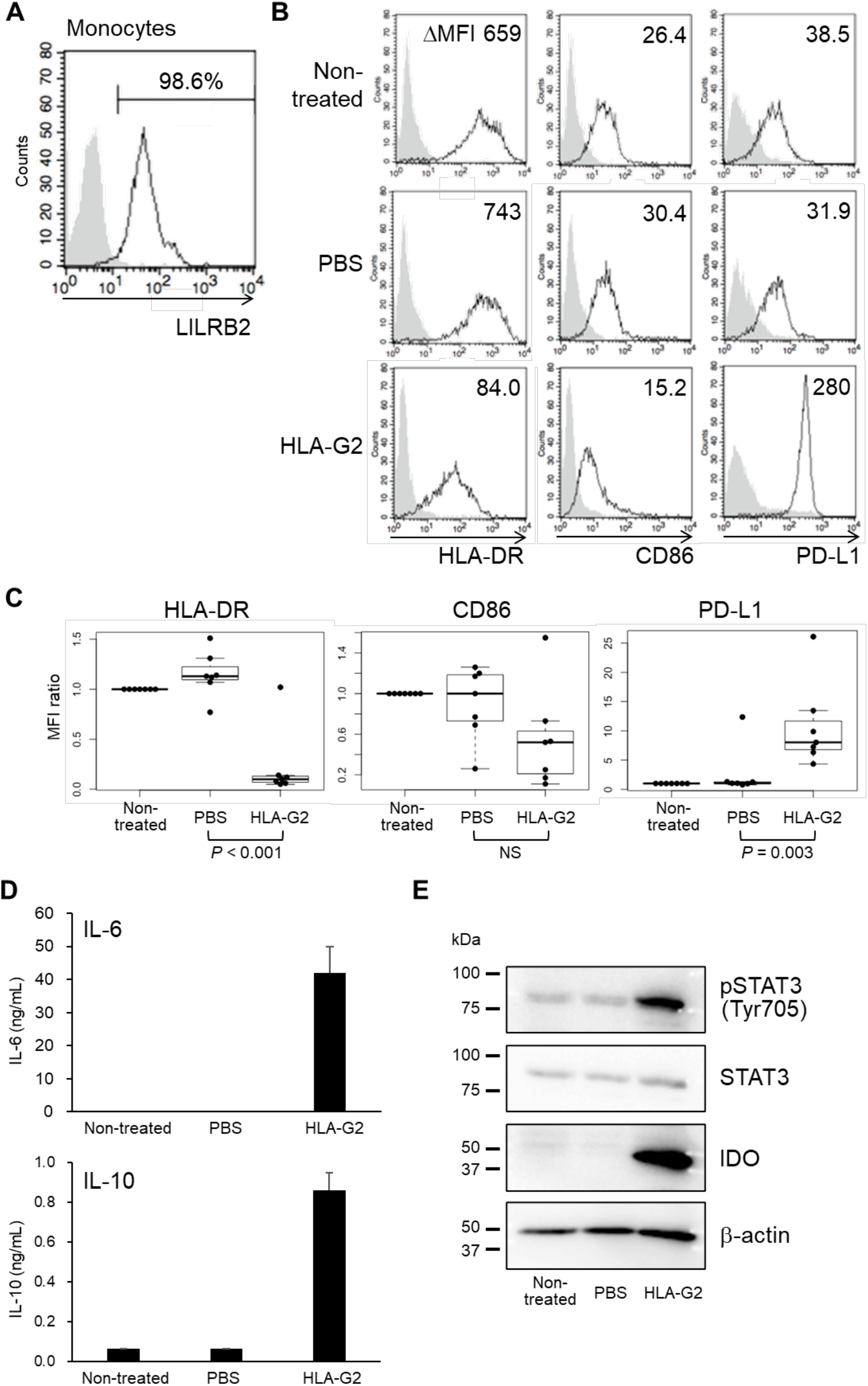
HLA-G2 induces human CD14-positive monocytes with an immunosuppressive phenotype. A. Flow cytometry analysis of surface LILRB2 expression by purified CD14^+^ monocytes (open histogram). Gray-filled histogram represents isotype control. Representative data of eight experiments from seven donors are shown. Cells were stained using the antibodies listed in **Supplementary Table 1.** B. Flow cytometry histogram analyses of HLA-DR, CD86 and PD-L1 expression (open histogram) of CD14^+^ monocytes either untreated or following treatment with PBS on HLA-G2 as indicated. Gray-filled histograms represent isotype control. Representative data of seven donors are shown. ΔMFI (MFI of target molecule – MFI of isotype-matched control) are shown in the upper right side. C. Box and superimposed dot plot showing the MFI ratio from (B) of CD14^+^ monocytes either untreated or in response to treatment with PBS or HLA-G2 as indicated. MFI ratio was calculated by (ΔMFI of PBS or HLA-G2)/ ΔMFI of non-treated control (n= 8). The boxes indicate the first and third quartiles. The bold line in the boxes indicates median values. *P* value was calculated by *t*-test. NS means not significant. D. IL-6 (upper) and IL-10 (lower) production in monocytes with/without HLA-G2 treatment. Representative data (average with standard deviation from two independent experiments) of three individuals are shown. E. STAT3 activation and IDO expression in response to HLA-G2 treatment. Representative data from three independent experiments are shown.

CD14-positive monocytes were cultured with the recombinant HLA-G2 protein (2.3 μM final concentration in PBS) or with PBS as described in Materials and Methods. The expression of HLA-DR and the co-stimulatory molecule CD86 was significantly down-regulated by incubation with HLA-G2 for 48 hours (**Figure 1B, C**). Notably, the immune checkpoint molecule, PD-L1, was up-regulated by HLA-G2 treatment (**Figure 1B, C**). Although upregulation of PD-L1 was clearly observed after treatment with HLA-G2 for 18 hours, changes in cell surface expression of HLA-DR and CD86 did not appear until a later time point (**Supplementary Figure 3A**).

In order to evaluate the molecular mechanism of HLA-G2 signaling, levels of the immunosuppressive cytokine, IL-10, and stimulation of IL-6/STAT3/IDO pathway were determined. Upregulation of IL-6 by monocytes was observed from 8 hours and increased with time and dose following HLA-G2 treatment (**Figure 1D, Supplementary Figure 3B-D**). Furthermore, the immunosuppressive cytokine, IL-10, was also upregulated by HLA-G2 (**Figure 1D, Supplementary Figure 3D**). Phosphorylation of STAT3 and expression of IDO-1 were also upregulated by HLA-G2 treatment (**Figure 1E**). Thus, HLA-G2 treatment affects similar signaling pathways to HLA-G1-LILRB2 (19, 21, 22).

### HLA-G2 binding to LILRB2 is blocked with anti-LILRB2 antibody

To determine whether the immunosuppressive phenotype induced by HLA-G2 could be mediated by binding to the HLA-G2 receptor, LILRB2, HLA-G2 binding to LILRB2 was first assayed in the presence of anti-LILRB2 antibody. Previously the anti-LILRΒ2 monoclonal antibody, 27D6 (Thermo Fisher Scientific), has been shown to block HLA-G1−LILRB2 binding (28). Thus, the ability of the 27D6 antibody to block HLA-G2−LILRB2 binding was analyzed by surface plasmon resonance (SPR) analysis (**Figure 2A**). The HLA-G2 binding response to LILRB2 covered by 27D6 antibody (black line) was dramatically reduced compared to the response to LILRB2 (gray line) (**Figure 2A**), indicating that 27D6 antibody blocks HLA-G2 binding to LILRB2. We then examined the expression profiles of signaling molecules in human monocyte and treated with HLA-G2. The 27D6 antibody blocked IL-6 and IL-10 upregulation, and IDO-1 expression induced in HLA-G2 treated monocytes (**Figure 2B, C**). Thus, induction of the immunosuppressive cytokine productions by HLA-G2 was mediated by HLA-G2 binding to LILRB2.

**Figure 2.**
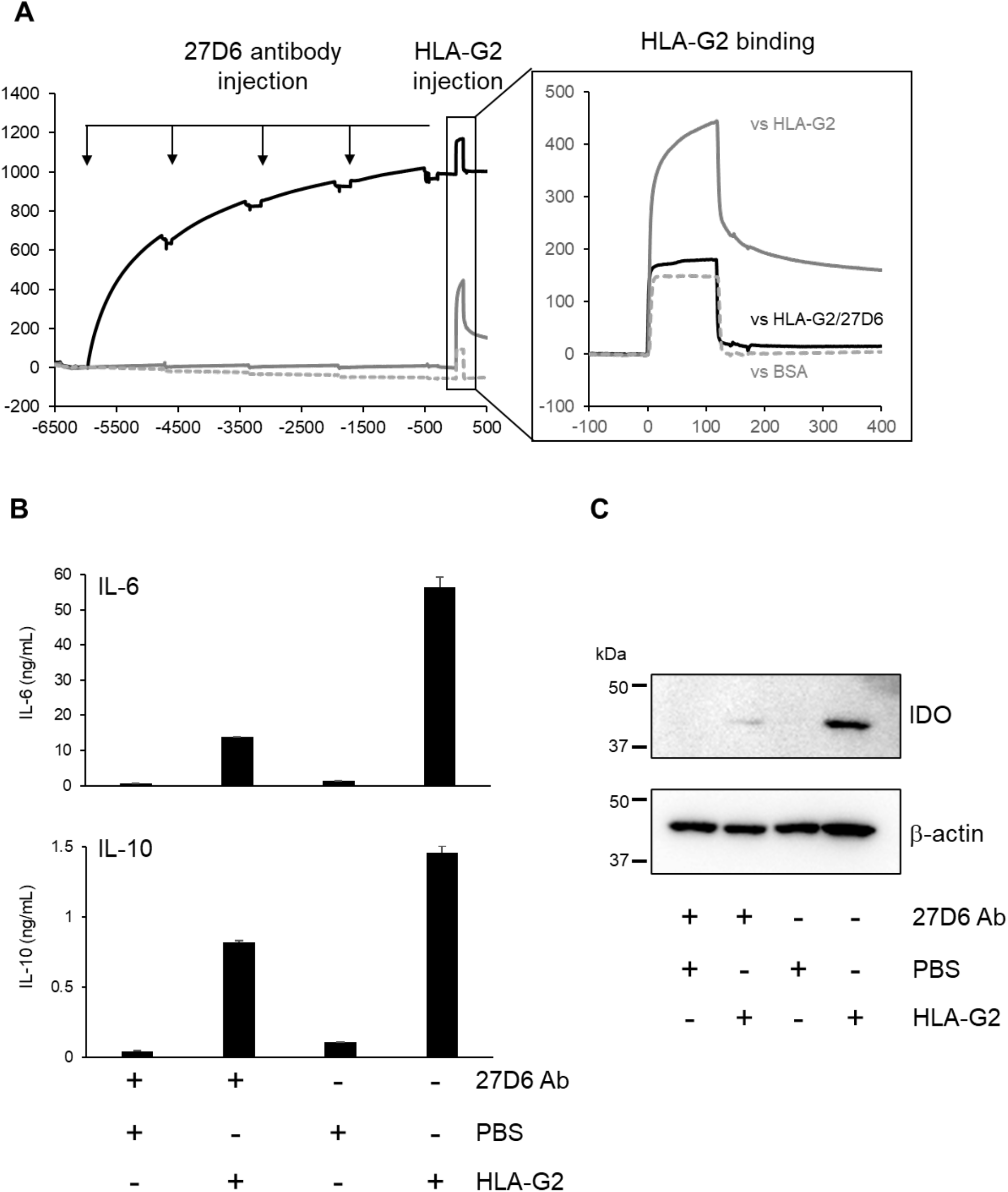
HLA-G2 binding to LILRB2 and induction of monocytes with an immunosuppressive phenotype are inhibited by LILRB2 antibody blockade. A. Left panel. SPR analysis of biotinylated BSA (control, gray dot line) or LILRB2 (black and gray solid lines) immobilized on a CAP chip via streptavidin (700 RU), and saturated with anti-LILRB2 (27D6) by repeated injection to cover the immobilized LILRB2 (black solid line). Right panel. SPR analyses showing binding responses of HLA-G2 against BSA (negative control), LILRB2 (positive control) and LILRB2 saturated with 27D6 antibody after Y-axis transformation at zero RU before injection. B. IL6 (upper) and IL-10 (lower) production by CD14^+^ monocytes in response to HLA-G2 treatment (2.3 μM) with or without 27D6 antibody. Representative data from three independent experiments using monocytes isolated from one individual. C. IDO expression by CD14^+^ monocytes in response to HLA-G2 treatment (2.3 μM) with or without 27D6 antibody. Representative data from three independent experiments.

### Induction of Tolerogenic IFN-DCs by HLA-G2−LILRB2 signaling

In order to determine whether HLA-G2 could also affect DC function, the levels of LILRB2 expression on PBMC-derived DCs were first determined. As previously reported (19), compared with monocytes (**Figure 1A)**, LILRB2 expression on IL-4-DCs was dramatically decreased following differentiation, while IFN-DCs maintained expression of LILRB2 (**Figure 3A**). Thus, the effect of HLA-G2 on DC phenotype was investigated using IFN-DCs. As observed with monocytes (**Figure 1**), HLA-G2 treatment also induced significant up-regulation of PD-L1 expression in IFN-DCs, but by contrast, HLA-DR and CD86 expression did not show any substantial change in expression (**Figure 3B**). Furthermore, HLA-G2 treatment enhanced production of IL-6 and IL-10 (**Figure 3C**).

**Figure 3.**
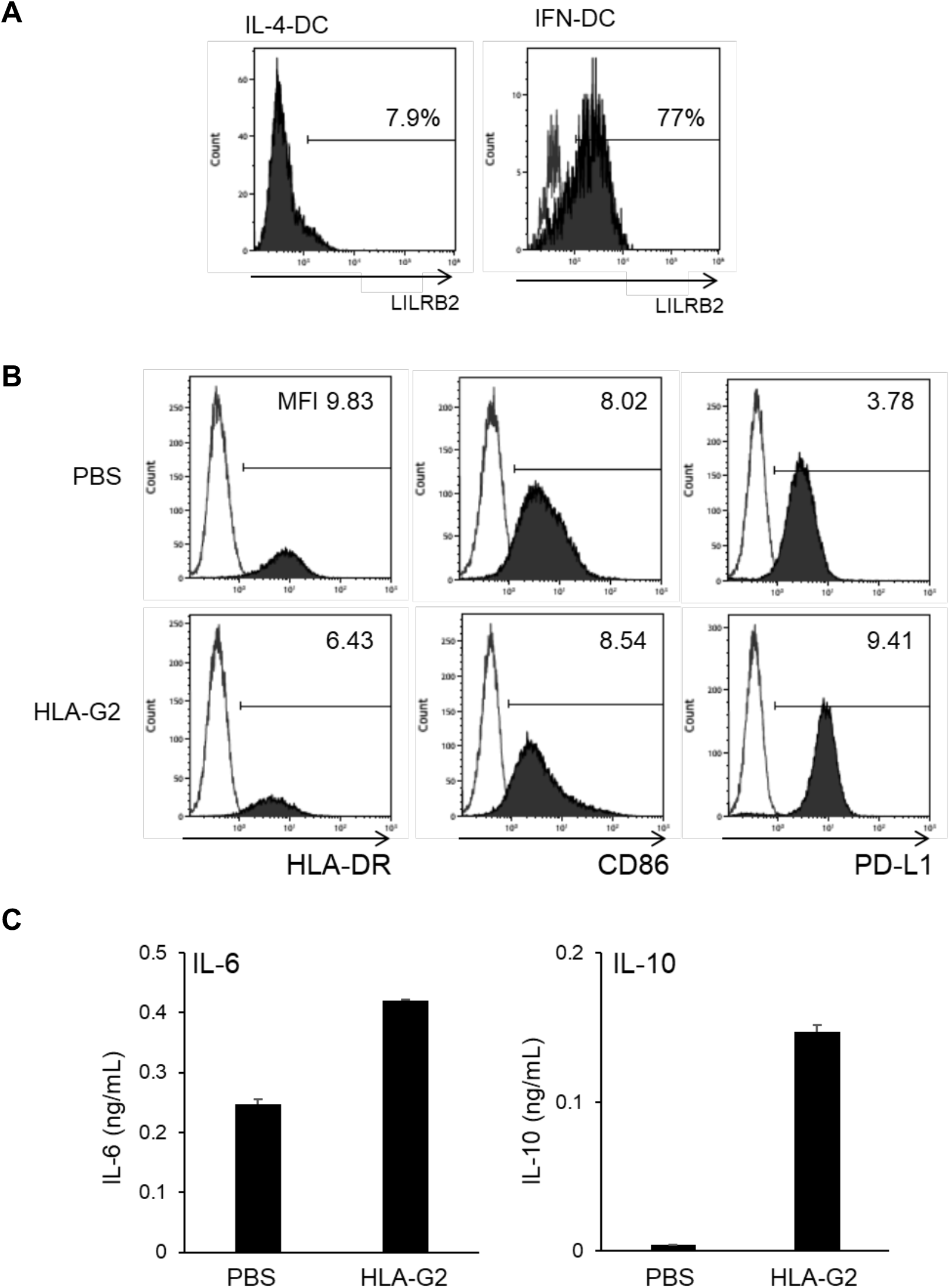
HLA_G2 induces IFN-Dendritic Cells (IFN-DC) with an immunosuppressive phenotype. A. Flow cytometry analysis of cell surface LILRB2 expression by IL-4 and IFN differentiated DC (black-filled histogram). Open histograms show staining with isotype control. Representative data from two individuals. Cells were stained using the antibodies listed in **Supplementary Table 1.** B. Flow cytometry histogram analyses of HLA-DR, CD86 and PD-L1 expression (black-filled histogram) by IFN-DC either untreated or treated with PBS or HLA-G2 as indicated. Open histograms show staining with isotype control. Representative data of three independent experiments from one individual. Cells were stained using the antibodies listed in **Supplementary Table 1.** C. IL-6 (left) and IL-10 (right) production by IFN-DCs with/without HLA-G2 (2.3 μM) treatment. Mean values with standard deviation from three independent experiments from one individual.

To examine the effect of HLA-G2 treatment on the function of IFN-DCs, mixed lymphocyte reaction (MLR) analyses were performed. IFN-DCs were first incubated with HLA-G2 or PBS for 48 hours, then cocultured with CFSE-labelled CD14-negative lymphocytes containing T cells from a different donor for further six days. The percentage of proliferating CD3-positive T cells incubated with HLA-G2-treated IFN-DCs (9.2%) was lower than that incubated with PBS-treated IFN-DCs (19.0%) (**Figure 4A**). Next, autologous MLR assays using a melanoma antigen peptide, MART-1(A27L), specific to HLA-A*0201 was performed. The MART-1(A27L) peptide-loaded HLA-G2-treated IFN-DC and CFSE-labelled lymphocytes from the same HLA-A:0201-positive donors were cocultured, and the population of the proliferating MART-1(A27L)-specific CD8^high^ T cells was compared. The CD8^high^ cells staining with MART-1(A27L) tetramer was decreased by co-culture with IFN-DCs (1.2%) treated with HLA-G2, compared to PBS-treated IFN-DCs (8.2%) (**Figure 4B**). These results preliminary suggested that HLA-G2-treated IFN-DCs are functionally tolerogenic and inhibit the proliferation of T lymphocytes.

**Figure 4.**
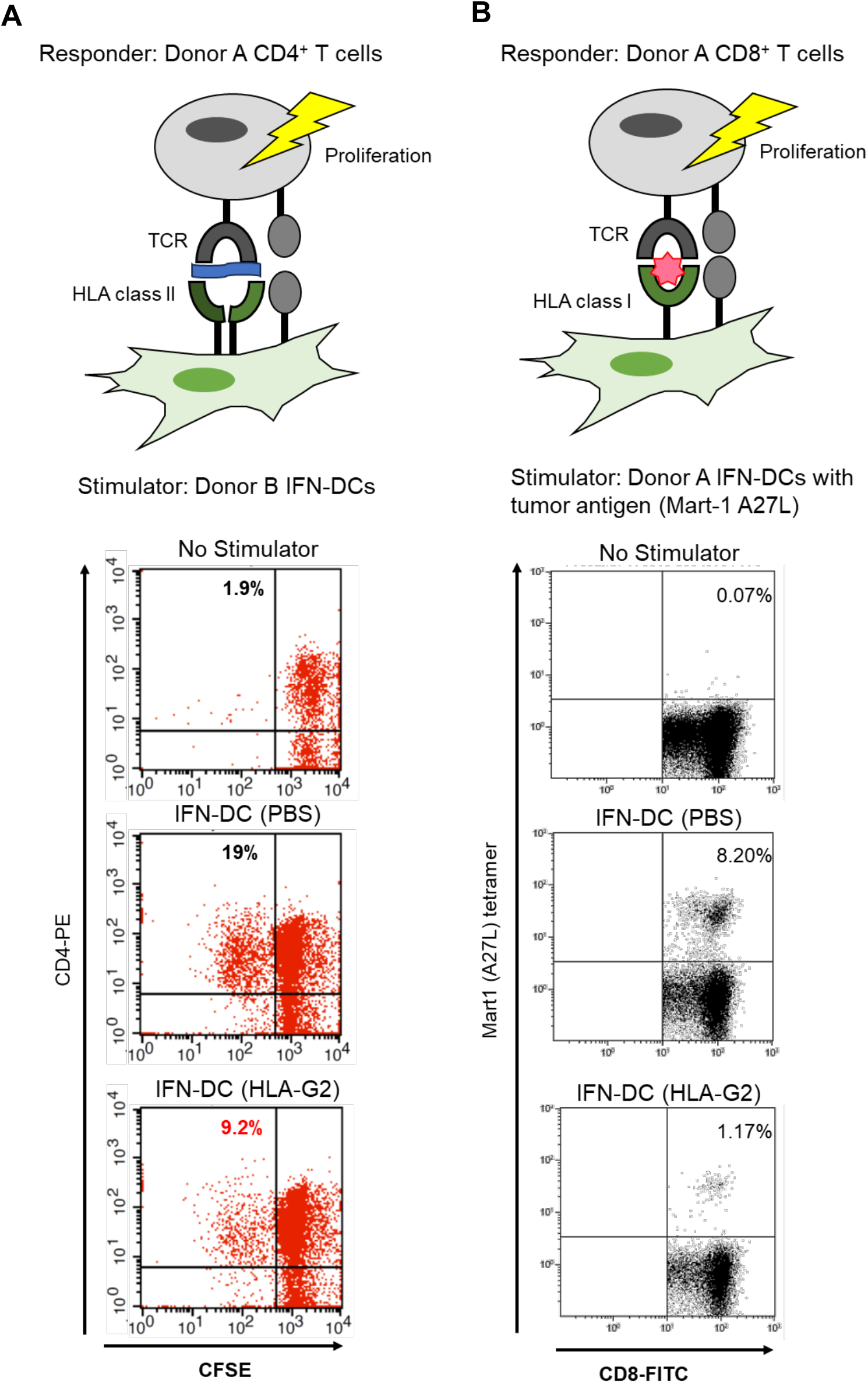
MLR of HLA-G2-treated IFN-DCs. A. Allogenic MLR analysis of CFSE-labelled T lymphocyte proliferation in response to IFN-DCs treated with or without HLA-G2 (2.3 μM) treatment. Dot plots show healthy donor CD3^+^ T cell proliferation after six days culture with treated or untreated IFN-DC and IL-2 from a different healthy donor. The percentage of CD4^+^CD3^+^ T cells are shown in the left upper panels. Representative data from two independent experiments using two different healthy donor pairs are shown. B. Autologous MLR analysis of CD8 T lymphocyte proliferation in response to culture with IL-2 and peptide-pulsed IFN-DCs from the same donor with or without HLA-G2 (2.3 μM) treatment. Dot blots show proliferation of CD8-positive T cells after 14-day stimulation. The percentage of CD8 positive T cells staining with MART-1 (A27L) tetramers cells are shown in the upper right quadrants. Plots show representative data from two independent experiments from one healthy HLA-A2-positive individual. Cells were stained using the antibodies listed in **Supplementary Table 1.**

## DISCUSSION

Here we demonstrated the immunosuppressive effects of HLA-G2 binding to LILRB2 expressed on human peripheral blood monocytes and monocyte-derived IFN-DCs. HLA-G2 has a much higher affinity to LILRB2 compared with other HLA class I molecules including HLA-G1 isoform due to dimerization (8). The phenotypes of HLA-G2 and the other HLA class I molecules via LILRB2 might be similar, but HLA-G2 is estimated to have a significant role in immune regulation via LILRB2 even at a lower concentration than other HLA class I molecules in vivo. Furthermore, since membrane-bound and soluble HLA-G2 share the same ectodomain solely responsible for receptor recognition, we especially focused on the function of the soluble HLA-G2 in regard to the possibility as a biopharmaceutical candidate targeting autoimmune diseases. To understand the molecular mechanism of immunosuppressive function induced by HLA-G2 treatment, we evaluated the expression profiles of representative cell surface molecules on human antigen presenting cells. HLA-G2 treatment down-regulated the expression of cell surface activation markers, CD86 and HLA-DR, on monocytes (**Figures 1B, C**). Notably, we observed substantial upregulation of PD-L1 on both monocytes and IFN-DCs (**Figures 1B, C, Figure 3B, Supplementary Figure 3A**). Upregulation of PD-L1 observed earlier than the downregulation of CD86 and HLA class II in IFN-DC means that PD-L1 signaling can modulate the expression of these molecules similar to the phenotype shown in GM-CSF-treated Bone marrow monocyte-derived DCs (29). Therefore, LILRB2 can modulate the maturation of DCs via regulation of the various surface markers’ expression. Since blocking the important PD-1/PD-L1 immune checkpoint pathway has shown promising results in clinical trials for various cancers (30), the observations in this study strongly suggest that the HLA-G2 binding to LILRB2 can be an important immune checkpoint pathway.

In regard to intracellular signaling, we showed that HLA-G2−LILRB2 interaction induced IL-6−STAT3−IDO-1 signaling in human antigen presenting cells (APCs) (**Figure 1**). While IL-6 has long been known as a potent proinflammatory cytokine in T cell activation (31), IL-6 can also have immunosuppressive functions, especially in the tumor microenvironment (32). Indeed, IL-6-STAT3 signaling inhibits DC maturation (33, 34) and IL-6−STAT3−IDO-1 signaling is activated in myeloid-derived suppressor cells (35). Furthermore, during DC differentiation, STAT3 is involved in the upregulation of PD-L1 in the differentiation of tolerogenic APCs (36). HLA-G2 binding to LILRB2 could be one mechanism by which IL-6 induces suppressive immune cells, as we showed with human monocytes and IFN-DCs in this study (**Figure 1D, Figure3C, Supplementary Figure 3B-D**). While it is unclear what types of DCs expressing LILRB2 are affected by HLA-G2 in vivo under physiological and pathological conditions, recently, LILRB1 and LILRB2, expressed on human dermal CD14-positive DCs, have been reported to inhibit the differentiation of naïve CD8-positive T cells into CTLs (37). Further analyses targeting other types of LILRB2 positive DCs and exploring unknown HLA-G2 receptors on immune cells should be analyzed in the future.

The induction of human myelomonocytic cells with an immunosuppressive phenotype by HLA-G2 treatment could be blocked with anti-LILRB2 antibody (**Figure 2**). The molecular mechanism of the immunosuppressive effect of LILRB2 ligation by HLA-G has only been reported using the most conventional HLA-G isoform, HLA-G1. In LILRB2 transgenic mice, HLA-G1 downregulated HLA-DR in DCs through the upregulation of IL-6 and IL-10 as well as STAT3 activation (21). Thus, we propose a potential molecular signaling network where HLA-G2 binding to LILRB2 leads to the induction of PD-L1 expression (**Figure 5**). The phosphorylated ITIMs of LILRB2 possibly recruit SHP-2 phosphatase (21). Subsequently, IL-6 and IL-10 would be up-regulated by NF-κB-related signaling to act in an autocrine manner as proposed in HLA-G1−LILRB2 signaling pathway model (21). Following this, STAT3 activation would lead to up-regulation of PD-L1, induction of IDO-1, and consecutive downregulation of HLA class II and CD86 (**Figure 5**). To further understand the molecular mechanisms of HLA-G2-induced immune suppression, a more detail functional relationship between immune checkpoint molecules, such as PD-L1-related molecules, needs to be elucidated.

**Figure 5.**
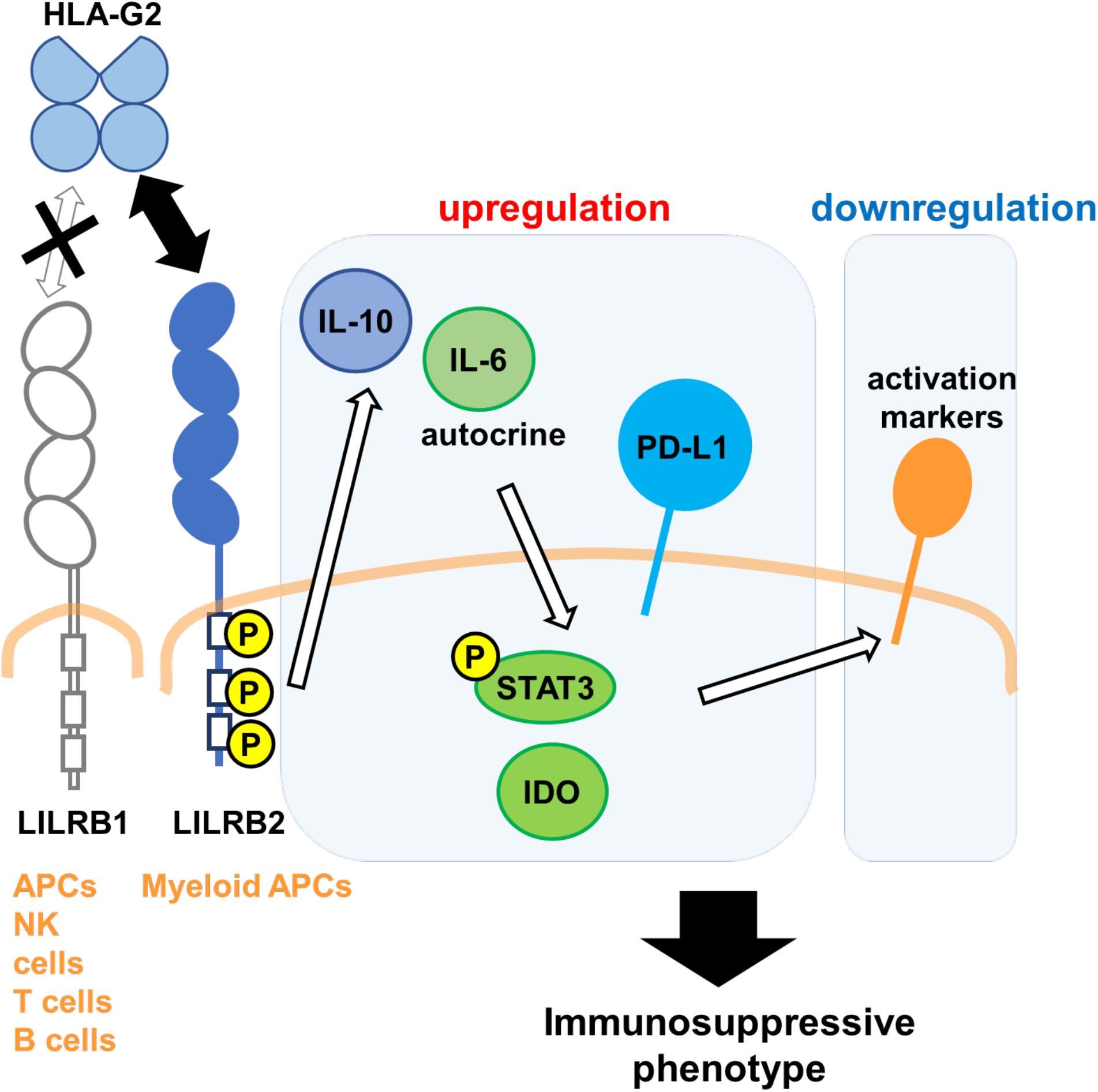
Model of the immunosuppressive mechanism of HLA-G2 in LILRB2-positive myelomonocytic cells.

Although the LILRB2 signaling by HLA-G1 and -G2 are similar as mentioned above, their receptor preference is clearly different. HLA-G1 can bind to both LILRB1 and LILRB2, however, HLA-G2 only binds to LILRB2. Thus, HLA-G2 functional effects are restricted to LILRB2-positive myeloid APCs but HLA-G1 can more broadly affect the function of LILRB1- and LILRB2-positive immune cells (APCs, B cells, NK cells and T cells). In mice, both HLA-G1 and HLA-G2 recombinant proteins bound to PIR-B, which is a mouse orthologue of LILRB2, and expressed on myeloid APCs and B cells. The immunosuppressive effects of HLA-Gs in mice immune inflammation models suggests that the direct suppression of T and NK cells is not essential to reduce autoimmune pathology (12, 16). When HLA-G2 is administered to humans, it binds only to LILRB2 expressed on APCs, so it likely induces more specific immune suppression than HLA-G1. Interestingly, the immunosuppressive effect in CIA mice was observed by the administration of less recombinant HLA-G2 protein than HLA-G1 and could possibly be due to higher affinity of HLA-G2 for PIR-B (11, 12). Regarding human LILRB2 binding, HLA-G2 showed higher affinity than HLA-G1 due to an avidity effect (8). In CIA mice, HLA-G2 showed significant immunosuppressive effects by less dose compared with the monomeric isoform, HLA-G1 (11, 12). Because the distribution of HLA-G2 receptor in human (LILRB2) and mice (PIR-B) is similar, the phenotype observed in vivo will be expected in human as well. Thus, in terms of the development of biopharmaceuticals for autoimmune disorders, HLA-G2 might be more suitable for clinical use in respect of the more restricted target specificity and higher receptor binding affinity.

HLA-G has recently been considered as a target for novel combined immunotherapies. Some tumors have been reported to express various soluble HLA-G molecules, such as β2m-free form of HLA-G1, the disulfide-bonded homodimer isoforms of HLA-G1, and HLA-G2 to suppress immune cell surveillance functions as summarized by Dias, F.C. et al (38). In epithelial cell adhesion molecule-positive colorectal cancer cells, not only PD-L1 but both LILRB2 and HLA-G were upregulated and associated with lymph node metastasis (39). Consistently, we showed that HLA-G2-treated IFN-DCs possibly inhibited T cell proliferation against both auto- and allo-antigens in MLR assays (**Figure 4**). Thus, our results raise the possibility that blocking HLA-G2−LILRB2 signaling could be an effective cancer treatment especially for the non- and low-responders to PD-1/PD-L1 therapy. Interestingly, the importance of PD-L1 expression in the accumulation of senescent cells and inflammation associated with aging was reported as a new aspect of PD-L1 signaling (40). The molecular relationship between HLA-G2−LILRB2−PD-L1 signaling and the regulation of expression of these molecules and inflammation in aging is worthy of future study.

In the present study, the number of donors available for the repeated analysis using fresh PBMCs and the optimal combination for the MLR assays are limited. We will clarify which molecules are dephosphorylated or regulated by the pathway activated by HLA-G2 through LILRB2 ITIMs and the profiles of each cell type expressing LILRB2 by HLA-G2 stimulation.

The present study showed the LILRB2 signaling by binding to HLA-G2 induced broad immune suppression and a tolerogenic phenotype in human CD14-positive monocytes and monocyte-derived IFN-DCs. These results suggest the functional importance of the HLA-G2 isoform and LILRB2 as immune checkpoint molecules. Thus, the HLA-G2−LILRB2 signaling pathway might be a promising target to regulate pathological immune responses in immune checkpoint-related diseases, such as autoimmune diseases and cancers. In summary, HLA-G2 modulates human immune responses by the induction of immunosuppressive monocytes and DCs with increased PD-L1 expression.

## Acknowledgement

We thank Y. Eiraku, T. Tadokoro for technical support and S. Kollnberger for helpful discussions.

## Author contributions

A.T., K.K., M.N., and K.M. designed the experiments, performed research, and analyzed the data. A.T., K.K., and K.M. wrote the paper.

## Data Availability

All data are included in the manuscript and its supplementary material.

