## Supplementary Figures 1-3 for "The human immune checkpoint molecule, HLA-G2, induces tolerance in monocytes and dendritic cells via upregulation of PD-L1"

### Takahashi et al., Supplementary Figure 1

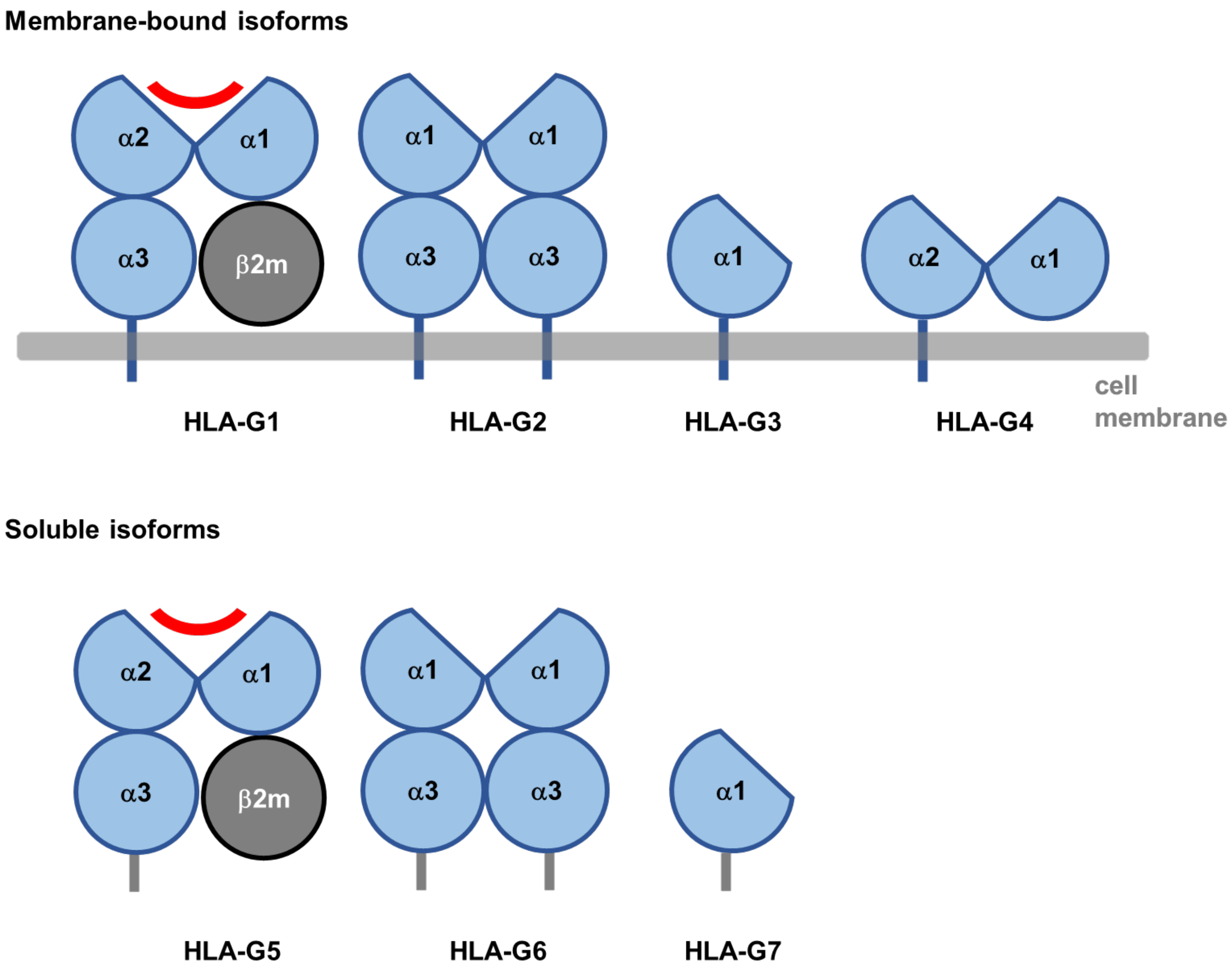


**Supplementary Figure 1. Schematic structures of HLA-G isoform proteins.**
The alternative splicing of the primary HLA-G transcript generates seven isoforms. HLA-G1 to HLA-G4 isoforms have a transmembrane region (membrane-bound forms), whereas HLA-G5 to HLA-G7 isoforms have an intron-encoded region at the C-terminus shown as a grey stalk region. In this study, HLA-G2 refers to HLA-G2 and HLA-G6 because these have the same ectodomains.

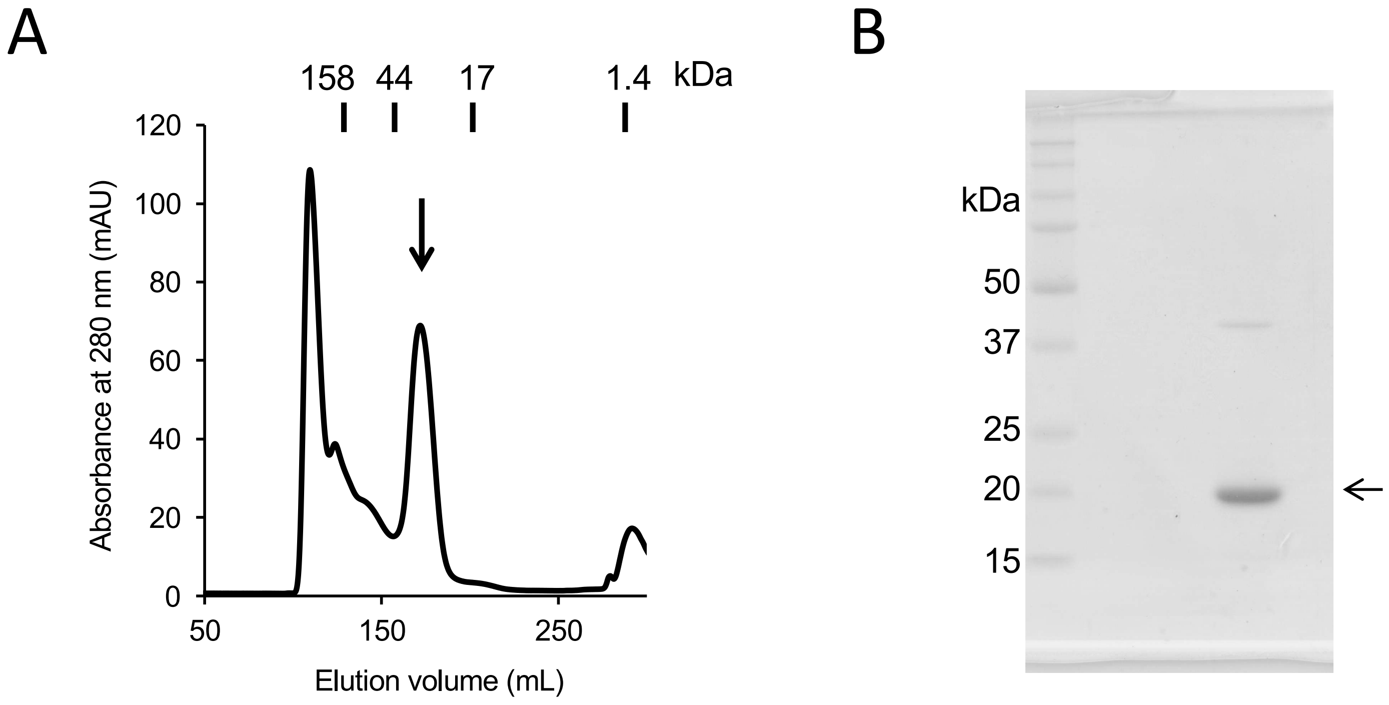


**Supplementary Figure 2. Preparation of HLA-G2 homodimer.**
A. HLA-G2 homodimer was purified by gel filtration chromatography on a HiLoad26/60 Superdex 75 column (42 kDa in dimer, left).
B. The peak indicated by a black arrow in A. was analyzed by SDS-PAGE under non-reducing condition (21 kDa in monomer). A faint band around 42 kDa indicates further oligomerization, such as a disulfide-linked dimer of HLA-G2 homodimer.

#### Takahashi et al., Supplementary Figure 3

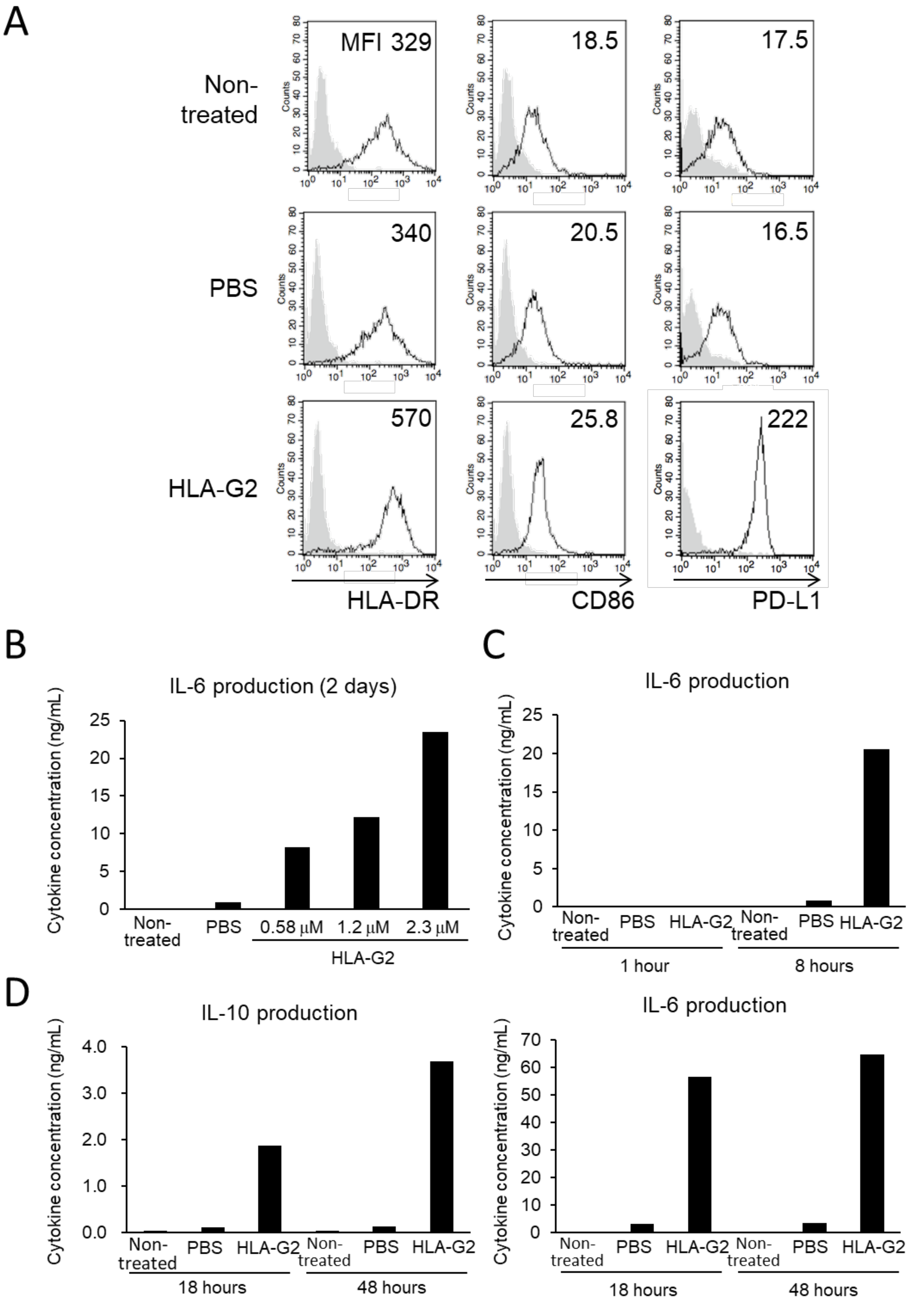


**Supplementary Figure 3. Time and dose-dependence of HLA-G2 induction of human CD14+ monocytes with an immunosuppressive phenotype.**

A. Flow cytometry histogram analyses of HLA-DR, CD86 and PD-L1 expression (open histogram) by CD14$^{+}$ positive monocytes in response to HLA-G2 treatment for 18 hours. Gray-filled histograms show isotype control antibody staining. Representative data from three independent experiments are shown. ΔMFI (MFI of target molecule – MFI of isotype-matched control) are shown in the upper right of the plots.

B. IL-6 production by HLA-G2 treatment (0.58, 1.2 and 2.3 μM) of CD14+ monocytes for 48 hours.

C. IL-6 production by HLA-G2 treatment (2.3 μM) of CD14$^{+}$ monocytes for one hour and 8 hours.

D. IL-10 (left) and IL-6 (right) production by HLA-G2 treatment (2.3 μM) of CD14$^{+}$ monocytes for 18 and 48 hours.

B-D. Data are representative data from duplicate experiments with CD14$^{+}$ monocytes from one individual.
